# Sociality shapes context-dependent stress responses to anthropogenic disturbance in African elephants

**DOI:** 10.64898/2026.09.30.755628

**Authors:** Kyla R. Funk, Levi Newediuk, Leslie R. Brown, Andre Ganswindt, Eric Vander Wal

## Abstract

Social relationships can buffer individuals against environmental challenges, reducing the physiological costs of disturbance. Assessing how social and environmental conditions jointly shape physiological stress is critical for predicting individual responses to disturbance, particularly in managed populations. We evaluated whether social and environmental stressors act additively or whether social context modifies physiological responses to environmental challenges. We combined behavioural data and faecal glucocorticoid metabolite samples from a small, intensively monitored population of African elephants (*Loxodonta africana*) in a fenced reserve. Using a Bayesian framework, we tested how social factors interacted with variation in the biophysical environment, including natural environmental and anthropogenic disturbances, to predict glucocorticoid concentrations. Natural environmental variables, including weather, water availability, and season, did not predict glucocorticoid concentrations. In contrast, helicopter use, a common management intervention, was associated with higher glucocorticoid concentrations. Social effects were context dependent. Following disturbance, low-ranking individuals showed higher glucocorticoid concentrations than high-ranking individuals, whereas this pattern reversed under low-disturbance conditions. Individuals with more social connections had lower glucocorticoid concentrations, but this effect was evident only under non-disturbance conditions. Measures of betweenness centrality did not predict glucocorticoid levels. These results demonstrate that social factors mediate physiological responses to anthropogenic disturbance and that management actions can impose uneven physiological costs across individuals. Quantifying interactions between social factors and the biophysical environment may improve conservation and management strategies for social species. Integrating physiological responses to interactions between social factors and the biophysical environment can guide conservation strategies and improve management interventions in social species.

**Lay summary:** Conservation actions can reshape the environments animals live in and the social systems they depend on, with consequences for well-being and survival. We show that in African elephants, human disturbance increases physiological stress, but social status determines who is most affected. Lower-ranking individuals show stronger stress responses after disturbance, whereas patterns reverse under non-disturbance conditions. Social connectedness can also reduce stress, although this effect varies across environmental conditions, highlighting how social factors and environment jointly shape physiological responses.

## 1. Introduction

Animals have multiple ways to cope with stressful environments. The physiological stress response is a series of adaptive mechanisms functioning in protecting an organism and restoring it to homeostasis (Ganswindt et al., 2010). Living in social groups can also help animals to cope with stress, as social partners can buffer the severity of perceived threats and consequently have an effect on the physiological coping mechanism (Kikusui et al., 2006; MacLeod et al., 2023). However, social groups can also increase stress for individuals of low rank, those who are poorly connected within the group, those facing intense social competition for resources, or individuals exposed to frequent aggressive interactions (MacLeod et al., 2023). Conversely, maintaining dominant social positions or extensive social relationships may itself incur physiological costs through increased social monitoring, vigilance behaviours, and relationship maintenance (Gesquiere et al., 2025; Milewski et al., 2022; Sapolsky, 2005). Group living can further be perceived as stressful when the social structure is unstable, when social isolation occurs within groups, or when individuals act as social buffers for others and incur their own physiological costs (Brandl et al., 2022; Kikusui et al., 2006). If unfavourable conditions endure, prolonged elevations in stress-related hormone levels can reduce fitness through immune suppression and impaired reproductive success (Möstl & Palme, 2002; Munck et al., 1984; Tilbrook, 2000). Therefore, the social mechanisms animals use to deal with stress can influence their susceptibility to stress. However, animals experience stressors within a biophysical environment that constrains and shapes their social environment (Webber et al., 2023). The biophysical environment and social environment, therefore, act together to shape the impact of the perceived stress, with each either compounding stress from the other or mitigating its effects (Dantzer et al., 2014). Such an interaction raises the possibility that when either the social or the biophysical environment is excessively stressful, reducing one source of stress might buffer individuals against the other.

Interrogating physiological stress responses of animals allows insight into the relative stress-related impact of different stimuli. During the physiological stress response in vertebrates, the hypothalamic-pituitary-adrenal axis (HPA axis) is activated, resulting in the secretion of multiple hormones, including glucocorticoids (GCs; Ganswindt et al., 2010; Sapolsky et al., 2000). Glucocorticoids are not synonymous with “stress”; rather, they represent one component of the physiological processes that help organisms respond to environmental and social challenges (MacDougall-Shackleton et al., 2019). Higher levels of GCs shift energy to coping with a stressor (MacDougall-Shackleton et al., 2019), and consequently, elevated GCs indicate the animal is perceiving a stressor in its environment (Bonier, Martin, et al., 2009; Bonier, Moore, et al., 2009; Crespi et al., 2013; MacDougall-Shackleton et al., 2019). There are a multitude of intrinsic and extrinsic stressors that affect the HPA axis, resulting in a temporary elevation of GC levels (Crespi et al., 2013). In poor biophysical environments, resource scarcity, extreme temperatures, drought, and seasonal declines in forage or water availability can require energetic trade-offs that elevate GCs to support immediate survival (Anderwald et al., 2021; Dezetter et al., 2022; Zimmer et al., 2020). Anthropogenic pressures such as habitat fragmentation, human presence, noise, aircraft activity, or active wildlife management may pose additional biophysical stressors with the potential to elevate GC levels, alter behaviour, and reduce access to key resources (Boyle et al., 2021; Kleist et al., 2018; Mirante et al., 2025). Identifying which biophysical stressors elevate GCs allows researchers and managers to assess when animals are coping with the environmental, social, and anthropogenic conditions they experience.

Within these ecological constraints, social systems represent a second major driver shaping GC variation (Crespi et al., 2013). Sociality can both buffer and exacerbate stress, depending on an individual’s position in the social environment. Dominance interactions, aggression, competition for mates or resources, and instability within social hierarchies can elevate GC levels (Creel, 2005; Edwards et al., 2013; Muller et al., 2021), whereas strong affiliative bonds and cohesive group structure can reduce GCs through social buffering (Wittig et al., 2016). Fission–fusion dynamics, dispersal events, and variation in social connectedness can further influence stress physiology by altering predictability of social interactions, risks of competition, and access to allies (Archie et al., 2006; Farine & Whitehead, 2015; Goldenberg et al., 2016; Wey et al., 2008). As individuals within the same landscape do not experience their social environment equivalently, variation in baseline GCs is expected to reflect both ecological pressures and social challenges. In social species specifically, such GC variation may carry substantial fitness consequences, as prolonged elevations in GCs can impair immune function, reproduction, and survival, while also influencing social behaviour that may serve potential buffering functions. Disruption of social structure through habitat loss, fragmentation, or management interventions may therefore amplify physiological stress by simultaneously increasing exposure to environmental challenges and weakening social buffering mechanisms. As a result, social animals experiencing human disruptions could face both direct (via habitat degradation, etc.) and indirect (via destabilized social structures) consequences, with disproportionately high GC-mediated costs. Monitoring baseline GCs, along with behavioural and other physiological indicators, can help identify costs and benefits as we gain insight into how animals are coping, or not coping, with environmental challenges and human interventions.

African elephants (*Loxodonta africana*) present a conservation paradox. Elephants are globally endangered as a species vulnerable to habitat loss and fragmentation, but are often locally hyperabundant and involved in human-wildlife conflict (Gobush et al., 2021; Pretorius et al., 2019; Thouless et al., 2016). As densely populated anthropogenic areas and agricultural areas are expanding across southern Africa, elephants are often confined to fenced reserves to minimize such conflict (Cumming & Jones, 2005). In South Africa, many elephant populations reside on small to medium-sized reserves, which limit movement and prevent migration. Such confined populations can reach high densities, and concerns over their potential impacts on ecosystem functioning and associated effects on habitats and other species have led to intense management interventions (Guldemond et al., 2017; Kerley & Landman, 2006). Management interventions for elephants can include fencing, translocation, contraception, water provisions, culling, or a combination of these strategies (Scholes & Mennell, 2008). Faecal glucocorticoid metabolites (fGCMs), which are excreted in dung, provide a useful non-invasive measure of adrenal glucocorticoid output over the preceding hours to days (Viljoen, Ganswindt, Palme, et al., 2008), which can be used to measure physiological stress of specific interventions or environmental conditions. Several studies have linked elevated fGCM concentrations to translocation, anthropogenic pressure, environmental conditions, and social dynamics in African elephants (Ganswindt et al., 2005; Oduor et al., 2024; Parker et al., 2022; Rasmussen et al., 2008; Viljoen, Ganswindt, Du Toit, et al., 2008). Elephants are highly social animals, living in complex, multi-tiered fission-fusion societies in which females form stable family groups that associate and separate with other family groups over time, while males disperse after adolescence and typically form fluid all-male groups, or roam independently (Archie et al., 2008; Wittemyer et al., 2009). Biophysical and social conditions are therefore likely to interact when shaping stress physiology, such that effects attributed to environmental stressors may partly reflect social context, and vice versa. Explicitly quantifying these interactions is particularly important for intensively managed endangered populations, where repeated human interventions may have cumulative physiological consequences that are unevenly distributed across individuals.

We tested two hypotheses concerning how biophysical and social environments shape individual variation in fGCM concentrations. Under the Additive Hypothesis (H1), biophysical and social challenges contribute independently (i.e., additively, without interaction) to physiological stress. We predicted:

- H1 P1: fGCM concentrations increase under challenging biophysical conditions or environments, including adverse weather, reduced water availability, and anthropogenic disturbance.
- H1 P2: fGCM concentrations will increase with social challenges such as higher rates of individual or intergroup aggressive interactions.

Under the Social Buffering Hypothesis (H2), social and biophysical environments interact to influence physiological stress through buffering effects (Cohen & Wills, 1985; Kikusui et al., 2006). We predicted:

- H2 P1: individuals embedded in a favourable social environment (having stronger social connections or higher dominance rank) will exhibit comparatively lower increases in fGCM concentrations in response to disturbances within their biophysical environments or other stressors compared to individuals within an unfavourable social environment.
- H2 P2: Under non-disturbance conditions, individuals with fewer social connections or lower dominance rank will exhibit comparatively lower fGCM concentrations, reflecting reduced physiological costs associated with maintaining dominant social positions or extensive social relationships when environmental constraints are minimal.

By integrating endocrine, behavioural, and social network data, we test how elephants physiologically respond to biophysical and management-related disturbances and social pressures (Table 1).

**Table 1.** Model structures and corresponding hypotheses.

| Model | Hypotheses and Predictions Tested | Independent Variable | Dependent Variables |
| --- | --- | --- | --- |
| Model 1 | H1 P1 | Individual fGCM concentrations from entire study period | -Rainfall<br>-Windspeed average<br>-Temperature average<br>-Water available<br>-Season<br>-Pressure<br>-Daily Temperature change |
| Model 2 | H1 P1 | Individual fGCM concentrations from entire study period | -Events on reserve |
| Model 3 | H1 P2 | Individual fGCM concentrations from entire study period | -Total aggressive interactions received<br>-Total aggressive interactions initiated<br>-Difference in aggressive interactions received and initiated |
| Model 4 | H1 P2 | Individual fGCM concentrations from entire study period | -Total daily intergroup aggressive interactions<br>-Total daily aggressions initiated by individual<br>-Total daily aggressions received by individual<br>-Total daily aggressions of individual (initiated + received) |
| Model 5 | H2 P1 | Individual fGCM concentrations subset to week following helicopter presence (07/26/2023-08/02/2023) | -Age<br>-Sex<br>-Degree centrality<br>-Betweenness centrality<br>-Dominance |
| Model 6 | H2 P2 | Individual fGCM concentrations subset to all data excluding the week following helicopter presence (01/01/2023-07/25/2023, 08/03/2023-04/01/2024) | -Age<br>-Sex<br>-Degree centrality<br>-Betweenness centrality<br>-Dominance |

fGCM: faecal glucocorticoid metabolite. Events on reserve category included: No events, hunting (of other species), others (additional people or vehicles), plane (taking off or landing on the property), helicopter (administering fertility control, aerial surveys).

## 2 Methods

### 2.1 Study site and animals

We conducted this study on Asante Sana Private Game Reserve (hereafter Asante Sana), Eastern Cape Province, South Africa (-32.29428, 24.97528; Clark & Vidal, 2019). The ∼11,000 hectare reserve comprises three main biomes: Albany Thicket, Grassland, and Nama-Karoo (Mucina et al., 2006). A boundary fence of 2.4 m height with 5 electrified strands surrounds the perimeter of Asante Sana, allowing for the passage of smaller vertebrate species, e.g. warthogs (*Phacochoerus africanus*), scrub hares (*Lepus saxatilis*), but is impermeable to larger species, e.g., elephants, zebra (*Equus burchellii*), buffalo (*Syncerus caffer*). The Milk River transects the study site, does not dam, and flows intermittently following sufficient rainfall. There are several human-made reservoirs throughout Asante Sana that are filled after rain but can dry up during prolonged droughts. The area receives an average of 516 mm of rain per year, with the wet season from October to April, and the dry season from May to September. The mean annual temperature is approximately 17°C, the mean minimum is 2°C, and the mean maximum is 34°C (Funk, Brown, et al., 2025). During winter months temperatures can fall as low as -4°C, where snow falls on the mountains, and in the summer, temperatures can exceed 40°C.

In 2004, a family unit of nine elephants (3 adult females, 4 juvenile females, and 2 calves) were introduced onto Asante Sana. In 2010, a single adult male was introduced, and from 2012 onwards, multiple calves were born. Eighteen elephants were present during the study period, including 7 adult females, 2 adult males, 4 juvenile females, and 5 juvenile males. There were no individuals experiencing energetically constraining life stages (i.e., lactating or pregnant females, weaning calves, males in musth) as male and female contraception has been used in the population since 2017. See population demographics in Table 2.

**Table 2.**
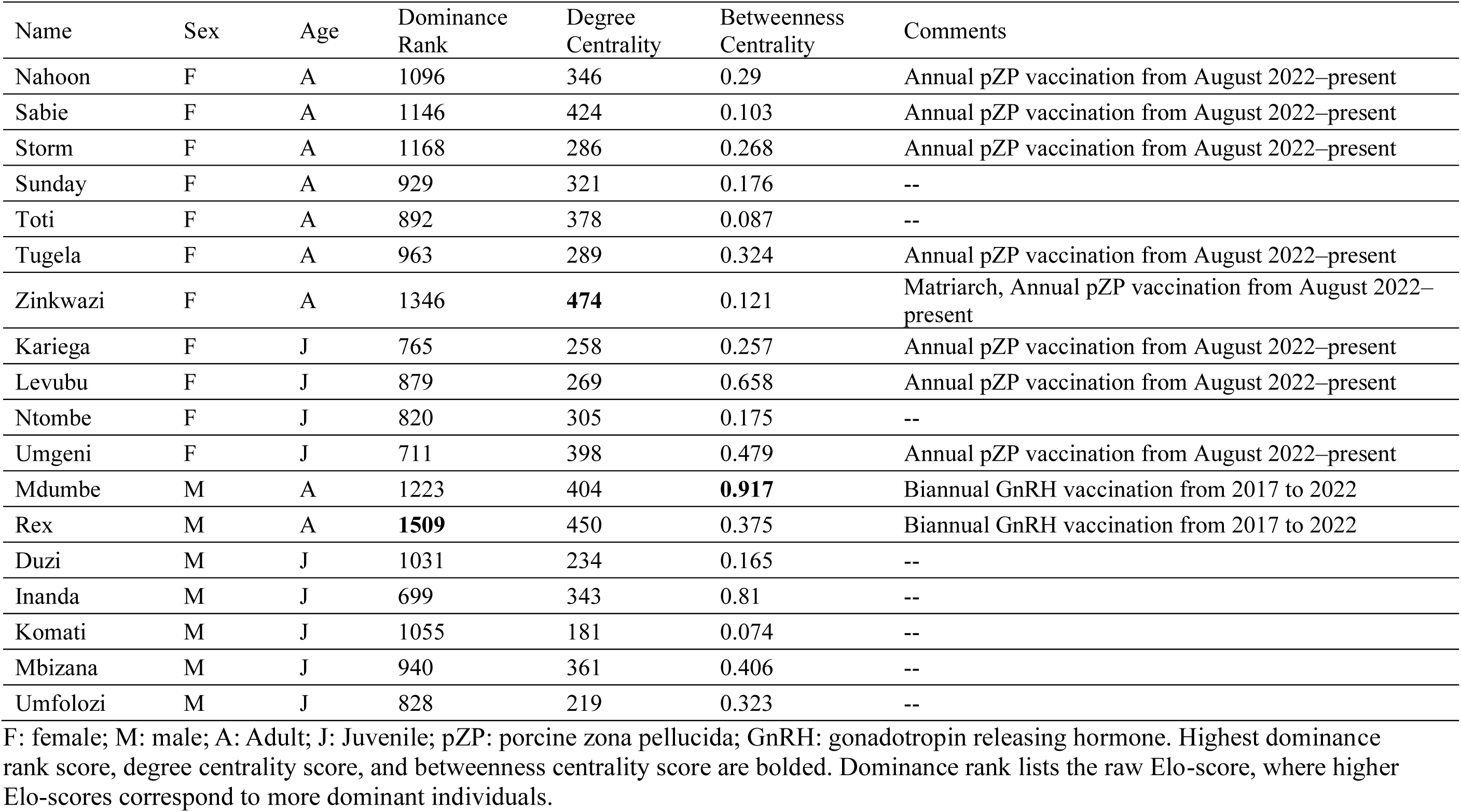
Study population of Asante Sana Game Reserve.

### 2.2 Data collection

Asante Sana is a private property that does not conduct ecotourism. As a result, the elephants of this study population were initially unhabituated to vehicles or people, as compared to habituated animals in South African National Parks or nature reserves. KRF began the process of habituating the elephants in July 2020. Habituation involved the repeated, non-invasive exposure of an animal to the researcher, research vehicle, and field equipment, until the individual no longer responded to these stimuli. Habituation is an essential fieldwork practice of behavioural field studies as it ensures that behavioural data reflect natural activity rather than the response to the observer and minimizes effects of an observer present. Behavioural data were therefore only collected once the elephants were habituated to the research team, and when no other people or vehicles (excluding the researcher and research vehicle) were present within a 300m radius of the elephant herd.

Between May 2020 and March 2024, we collected baseline data from individual animals, identified from their distinct ear notches and tusk shapes. We collected data within daylight hours at least 5 days per week, which consisted of scan samples (Altmann, 1974) collected on all visible animals every 30 minutes. We recorded the animal’s identity, behaviour, and distance and identity of their nearest neighbours. Every day of the study period, we also collected reserve-level information on variables of interest such as the water available, number of humans and vehicles present on the reserve that day, if a plane was active (taking off or landing on the property), if a helicopter was employed for contraception administration, or if there was any hunting of other species within the reserve that day. Weather data were collected using a Logia 5-in-1 weather station (model # LOWSC510WB) to record rainfall and other weather-related data such as temperature, windspeed, and atmospheric pressure.

We collected faecal samples between January 2023 and April 2024, spanning both the wet (October-April) and dry (May-September) seasons. Faecal samples were collected *ad libidum* from observed animals while collecting the baseline data described above. When the animal moved a sufficient distance away from the sample, approximately 20 grams (golf ball size) of well-mixed wet faeces were collected from the centre of the bolus to avoid cross-contamination with soil, water, or urine. Samples were placed and sealed in labelled plastic Ziplock bags. Samples were placed on ice immediately after collection and placed in a freezer at the end of each field day. Every sample was frozen within 8 hours of collection, and remained frozen during storage and transport, until analyses at the Endocrine Research Laboratory of the University of Pretoria, South Africa.

### 2.3 Extraction of faecal glucocorticoid metabolites

Faecal samples were lyophilized, pulverized and sifted using a nylon mesh strainer to separate undigested fibrous material from the faecal powder (Fieß et al., 1999; Ganswindt et al., 2005). Faecal GCMs were quantified using an Oxoaetiocholanolone enzyme immunoassay (measuring fGCMs with a 5β-3α-ol-11-one structure), as first described by Möstl et al., (2002), and validated for measuring adrenocortical activity in African elephants (Ganswindt et al., 2003, 2010). Assay sensitivity was 1.5 ng/g dry faecal mass, and intra-assay coefficients of variance, determined by repeated measurements of high- and low-value quality controls, were 5.57% and 6.58%, respectively, and the inter-assay coefficients of variance were 9.69% and 12.84%, respectively.

### 2.4 Social Network Measures

We constructed an undirected edge list using proximity data collected during baseline data collection. To determine network centrality of each individual, we calculated both degree centrality and betweenness centrality using the ‘igraph’ and ‘sna’ packages (Butts, 2024; Csárdi et al., 2025). Degree centrality is a measure of the number of connections a node (individual) has within the network, where betweenness centrality measures the extent to which a node lies between other nodes in the network and can be computed as the proportion of shortest paths that pass through the node (Tabassum et al., 2018).

### 2.5 Dominance interactions and calculating dominance hierarchy

We recorded all observed dominance interactions and identified the aggressor (the individual that initiated the aggressive interaction), the receiver (the individual that received the aggressive interaction) and the outcome of the interaction from the perspective of the aggressor (win, lose, draw, unknown). Dominance interactions included all overt aggressive and submissive interactions that were physical (chases, pokes, pushes) or nonphysical (supplants, displaces) following the protocol of Wittemyer & Getz (2007). Operational definitions of the dominance interactions observed are listed in Table A1. Using the temporal sequence of dominance interactions over the study period, we calculated individual dominance ranks as Elo-ratings (Albers & De Vries, 2001) with the ‘EloRating’ package in R (Neumann et al., 2011). The number of points allotted to an individual after an outcome is determined by the k value, which we set to the default value of 100 (Neumann et al., 2011). Higher Elo-scores correspond to more dominant individuals (winning more outcomes) and scores decrease as individuals are more subordinate (losing more outcomes). We then used the ‘ani-Dom’ package to calculate the repeatability score of ranks from randomisations of the interaction orders estimate the uncertainty of the dominance hierarchy, where a score of score above 0.8 suggests a reasonably robust hierarchy (Farine & Sanchez-Tojar, 2021). We received a repeatability score of 0.947, indicating a high level of certainty of our input data and stable dominance hierarchy. We constructed pairwise plots and calculated correlation coefficients for dominance, degree centrality, and betweenness centrality to assess collinearity between these variables (supplementary materials, Figure S1).

### 2.5 Statistical analysis

We analysed the data in a Bayesian framework, using RStudio version 2026.01.0+392 (Posit Team, 2025). To determine which predictor variables influenced fGCM levels, we generated six hierarchical models using the ‘brms’ package (Bürkner, 2017), with individual fGCM levels as the response variable. Predictor variables examined in each model are summarized in Table 1. Gastrointestinal passage of time in African elephants is generally 24-48 hours, with fGCM levels typically peaking 36 h after ACTH stimulation (Ganswindt et al., 2003; Manning et al., 2022). Consequently, all environmental, anthropogenic, and behavioural predictor variables were matched with fGCM samples collected 2 days after the observed event, ensuring the analyses reflect the biologically appropriate delay between exposure and fGCM excretion (Manning et al., 2022; Newediuk et al., 2026). Dependent variables and corresponding hypotheses and predictions for each model are listed in Table 1. Individual identities were entered as crossed random effects in all models. We set weakly informative priors centred on zero for all models, i.e. normal (0,1). Age and sex were included in models examining social network measures given some studies indicate these demographics influence GCs (Pokharel & Brown, 2023). We set adults and females as the reference levels for the age and sex categories, respectively. Numeric predictor variables were centered and scaled. We accounted for variation in the number of dung samples per individual by setting the total number of samples per individual as an offset variable in every model. We ran four chains for 2500 iterations for each model and confirmed convergence of the chains, i.e. all R-hat ≤ 1.01. Bulk effective sample size estimates (Bulk-ESS) and tail effective sample size estimates (Tail-ESS) confirmed that the posterior means and medians were reliable.

Our response variable of interest, fGCM level, was not normally distributed (supplementary materials, Figure S2), and so when fitting the models we specified a “skew normal” distributions. We used the posterior predictive check (‘pp_check’) function to assess the adequacy of model performance (Gabry et al., 2019) and the ‘testdispersion’ and “plotResiduals” functions of the ‘DHARMa’ package to test for dispersion issues (Hartig, 2017). We detected no dispersion issues in any of the models (supplementary materials, Figure S3). Model fit was assessed using the ‘loo_R2’ function of the ‘brms’ package, which computes a leave-one-out-(LOO)-adjusted R-squared for regression models (Bürkner, 2017). However, when comparing the Loo R2 value for each model, Model 5 initially had a negative R2 value, indicating a poor model fit, a common issue in Bayesian models (Gelman et al., 2019). All other measures of model performance were adequate for Model 5, however, this model did have less data than all other models (N = 49 observations). Outliers were noted, and so, we specified a “Weibull” distribution in order to model extreme values (Bürkner, 2017) as we did not want to remove data/outliers given the already small sample size. The new Weibull model had an adequate Loo R2 value and all other measures of model performance (supplementary materials, Figure S3, Table S1). The Weibull model was quantitively similar to the initial model (see supplementary materials), and we report and compare the findings from the original “skew-normal” fitted model (appendix A6).

We generated posterior density distributions and other graphical outputs with the ‘ggplot2’ package (Wickham, 2016), and the ‘bayestestR’ package (Makowski et al., 2019). We summarized model estimates with 95% credible intervals (CI). Effect certainty was quantified using the probability of direction (pd), which reflects the proportion of posterior samples supporting a positive or negative effect (Makowski et al., 2019). Thresholds of ∼97.5%, ∼99.5%, and ∼99.95% correspond to weak, moderate, and strong evidence, respectively (Colquhoun, 2014). To compare and visualize differences in model-predicted fGCM concentrations and social network measures, we extracted the posterior density distributions from rank, degree centrality, and betweenness centrality from each model, and split individuals into “low” and “high” categories for each predictor variable of each model, while treating all predictors as continuous in the statistical analyses (e.g., Nord et al., 2022).

## 3 Results

We collected 372 fGCM samples over the study period. Adult females had an average fGCM concentration of 0.375 (range = 0.137-0.814, sd = 0.135), adult males an average concentration of 0.358 (range = 0.182 – 0.711, sd = 0.111), juvenile females an average concentration of 0.361 (range = 0.118 – 0.807, sd = 0.133), and juvenile males an average concentration of 0.333 (range = 0.128 – 0.724, sd = 0.114). Main effects for posterior estimates, standard errors (SE), upper and lower 95% CIs, bulk and tail effective sample size estimates and probability of direction estimates are presented for all models in the appendix (Table A2-A7).

### 3.1 Biophysical Environment

When testing our first prediction under Hypothesis 1 (H1 P1), Model 1 indicated that weather related predictor variables such as rainfall (pd=97.94%), windspeed average (pd=64.98%), temperature average (pd=87.90%), water available (pd=50.22%), specific seasons (pd=91.62%, pd = 51.52%), atmospheric pressure (pd=75.04%), and daily change in temperature (pd=98.06%) did not predict fGCM levels. Although pd scores for rainfall and temperature change indicated effect certainty, the estimate for both rainfall (0.01) and temperature change (-0.01) were nearly 0, indicating there is a high certainty of no effect. However, when looking at events related to human activity in Model 2, helicopter use predicted higher fGCM levels (pd=99.38%). Other predictor variables of Model 2 including hunting (pd = 67.14%), other people/vehicle presence (pd=85.68%), and planes taking off or landing (pd = 83.68%), were not predictive of fGCM scores. We plotted the posterior expectations from Model 2 to examine the relationship between all human activity categories and fGCM levels, highlighting fGCM scores in relation to helicopter use (Figure 1).

**Figure 1.**
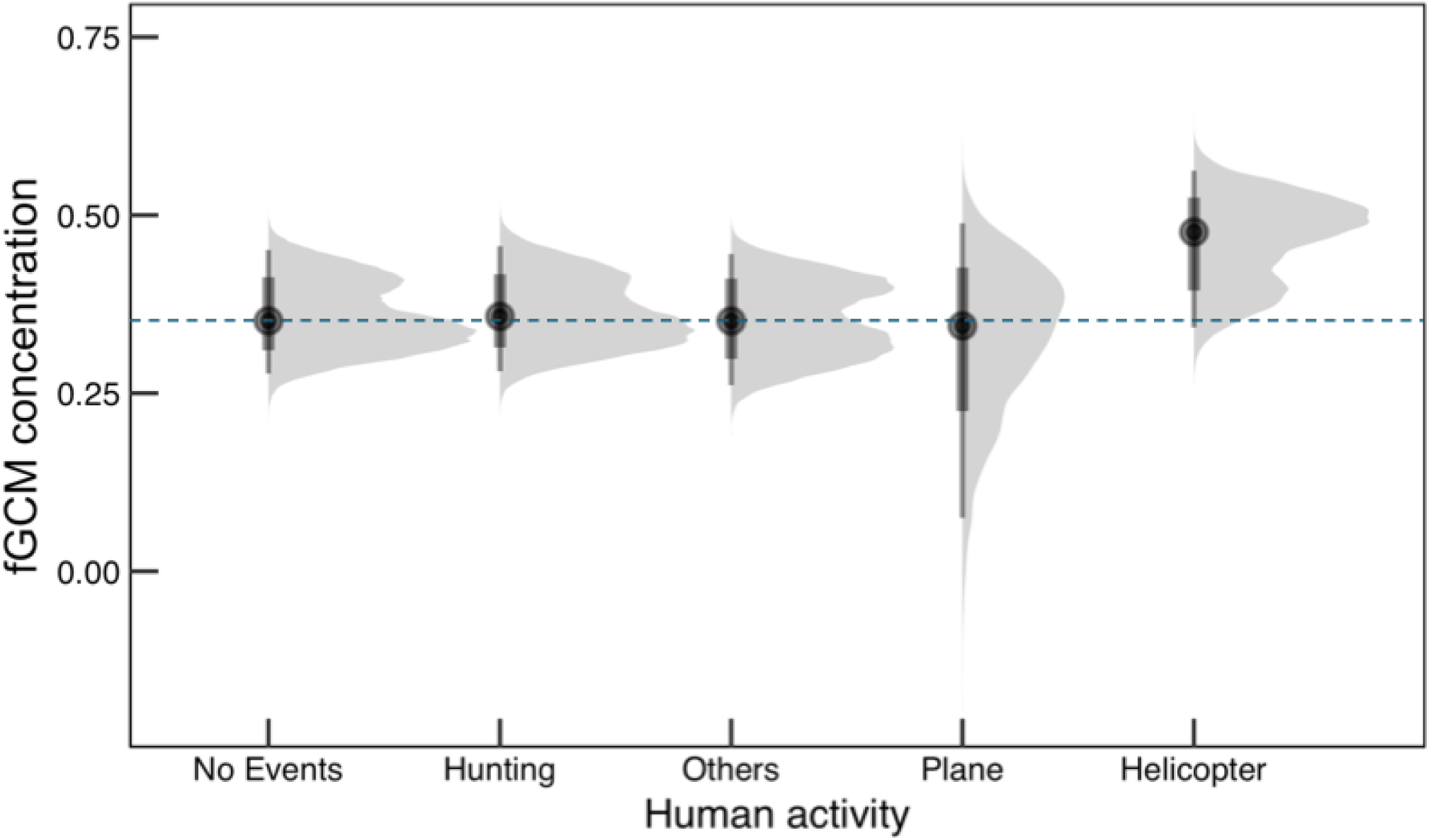
Posterior expectations extracted from Model 2 displaying the relationship between various human activities and faecal glucocorticoid metabolite concentrations (fGCMs). fGCM concentrations are in μg/g dry weight. Density plots indicate the range of fGCM levels predicted by the model for each variable of interest. The black dot represents the mean, and the dark bar represents the 95% credible intervals. We set our reference variable to days with “no events” and have a dashed line through the mean of this reference variable. Categories to compare include: hunting (of other species within the reserve), others (additional people or vehicles on the reserve), plane (taking off or landing on the property), and helicopter (administering fertility control, aerial surveys).

### 3.2 Social Environment

Aggressive interactions were not predictive of fGCM levels. In Model 3 (H1 P2), where total aggressive interactions over the study period were examined, the total aggressive interactions initiated (pd = 58.74%), total aggressive interactions received (pd = 74.98%), and the difference in aggressive interactions received and initiated by each individual over the study period (pd = 52.96%) were not predictive of fGCM levels. Daily aggressive interactions were also not predictive of fGCM concentrations, as indicated in Model 4 (H1 P2). Daily aggressive interactions initiated by an individual (pd = 50.02%), daily aggressive interactions received by an individual (pd = 51.06%), and total daily aggressions for an individual (pd = 50.66%) had pd scores that indicated no effect. Daily intergroup aggressive interactions had a pd score that indicated weak evidence in certainty for the direction of an effect (pd = 98.64%); however, the estimate for the effect was -0.02 indicating a very small or no effect, given the standard error of the estimate was 0.01 (Appendix A5).

### 3.3 Buffering effects of the biophysical and social environment

For Model 5, we investigated whether individual levels of social variables were predictive of fGCM scores in a stressful biophysical environment (H2 P1). Models 1–4 identified helicopter disturbance as the only anthropogenic variable associated with higher fGCM concentrations. Consequently, Model 5 examined social predictors during the week following helicopter activity (07/26/2023-08/02/2023), whereas Model 6 examined the remaining non-disturbance period (01/01/2023-07/25/2023, 08/03/2023-04/01/2024). However, for Model 5, degree centrality (pd = 93.36%), betweenness centrality (pd = 92.32%), and dominance rank (pd = 84.16%) were not found to be related to fGCM concentrations. In Model 6 we kept all of the same predictor variables, but excluded data following helicopter use to represent a non-disturbance period within the biophysical environment (H2 P2). Degree centrality score was found to be weakly predictive of fGCM scores, with higher degree centrality scores being predictive of lower fGCM concentrations (pd = 98.26%). Neither betweenness centrality (pd = 96.42%) nor dominance rank (pd = 86.70%) predicted fGCM concentrations.

During the helicopter disturbance period (Model 5), individuals with a low rank were more strongly associated with having a higher fGCM concentrations compared to individuals of high rank (Figure 2A). During non-disturbance periods (Model 6), the relationship was reversed, and high-ranking individuals were more strongly associated with having a higher fGCM concentrations than individuals of low rank. Individuals with a high degree centrality score were more strongly associated with lower fGCM concentrations in both non-disturbance periods, and disturbance periods following helicopter use (Figure 2B). Individuals with a high betweenness centrality score were more strongly associated with higher fGCM concentrations in both non-disturbance periods, and disturbance periods following helicopter use compared to individuals with a low betweenness centrality score. (Figure 2C).

**Figure 2.**
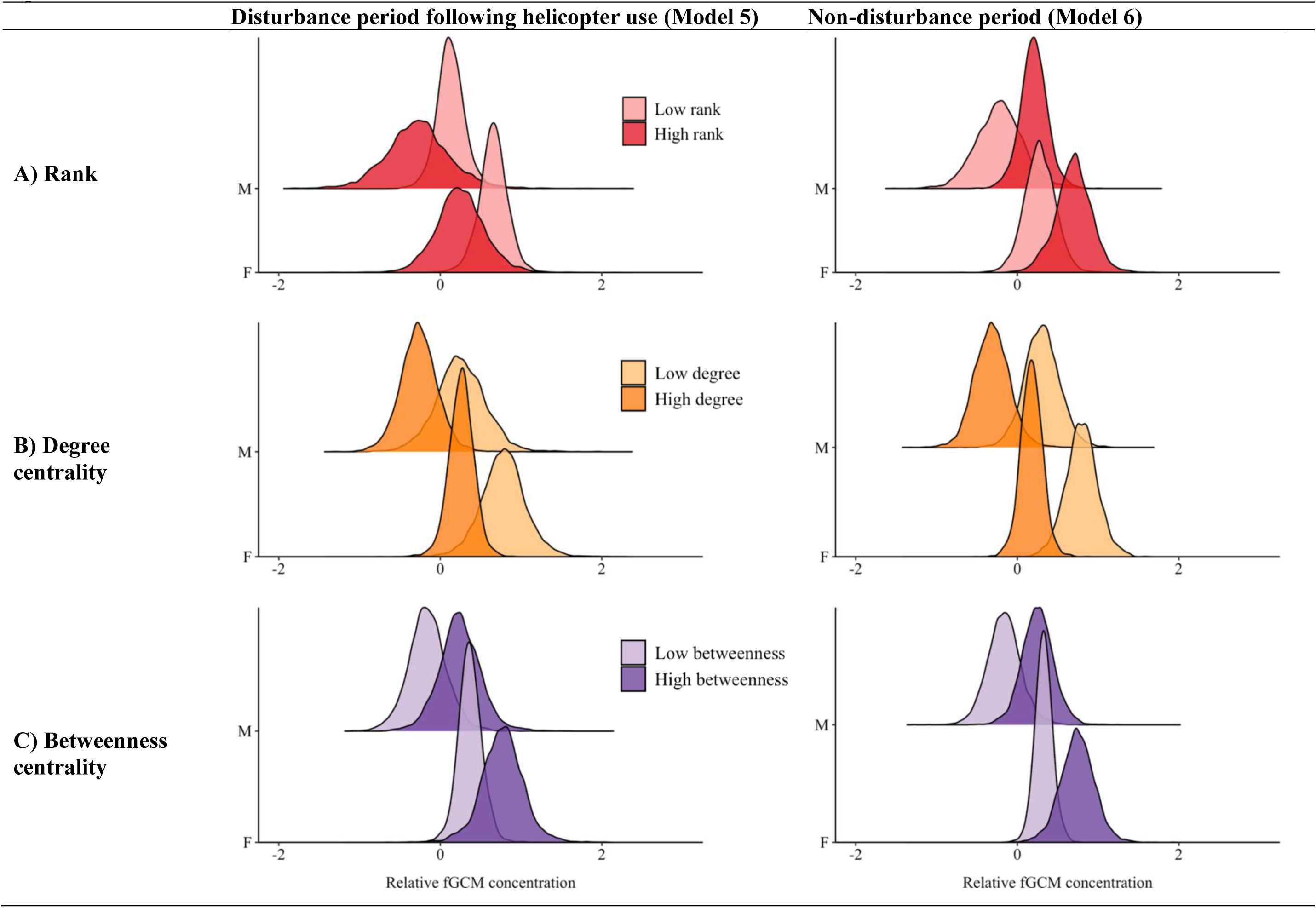
Relative fGCM concentrations predicted by model 5 and 6 for individuals with A) low and high dominance ranks, B) low and high degree centrality scores, and C) low and high betweenness centrality scores. “Disturbance period following helicopter use” indicates the period that elicited elevated fGCM concentrations (helicopter use, 07/26/2023-08/02/2023), and “Non-disturbance period” refers to the period that did not elicit heightened fGCM concentrations (helicopter data excluded, 01/01/2023-07/25/2023, 08/03/2023-04/01/2024). fGCM: faecal glucocorticoid metabolite. The height of the density curve indicates the probability of the predicted rate from the model, and the spread of the curve represents its uncertainty. F: female; M: male.

### 3.4 Age, sex, and group level effects

Sex predicted fGCM concentrations in both disturbance and non-disturbance periods, with males exhibiting lower fGCM concentrations than females in both contexts (Model 5: pd = 99.46%; Model 6: pd = 99.64%; Figure 3). However, the difference between the sexes was more pronounced during the disturbance period following helicopter activity. Age did not predict fGCM concentrations in Model 5 (pd = 95.24%) or Model 6 (pd = 96.34%). In all models, there was variation with respect to group-level effects across individual elephants (Table A2-A7), indicating that each elephant responded differently to the various predictors. We plotted the posterior expectations to depict the variation between each elephant in Figure 4.

**Figure 3.**
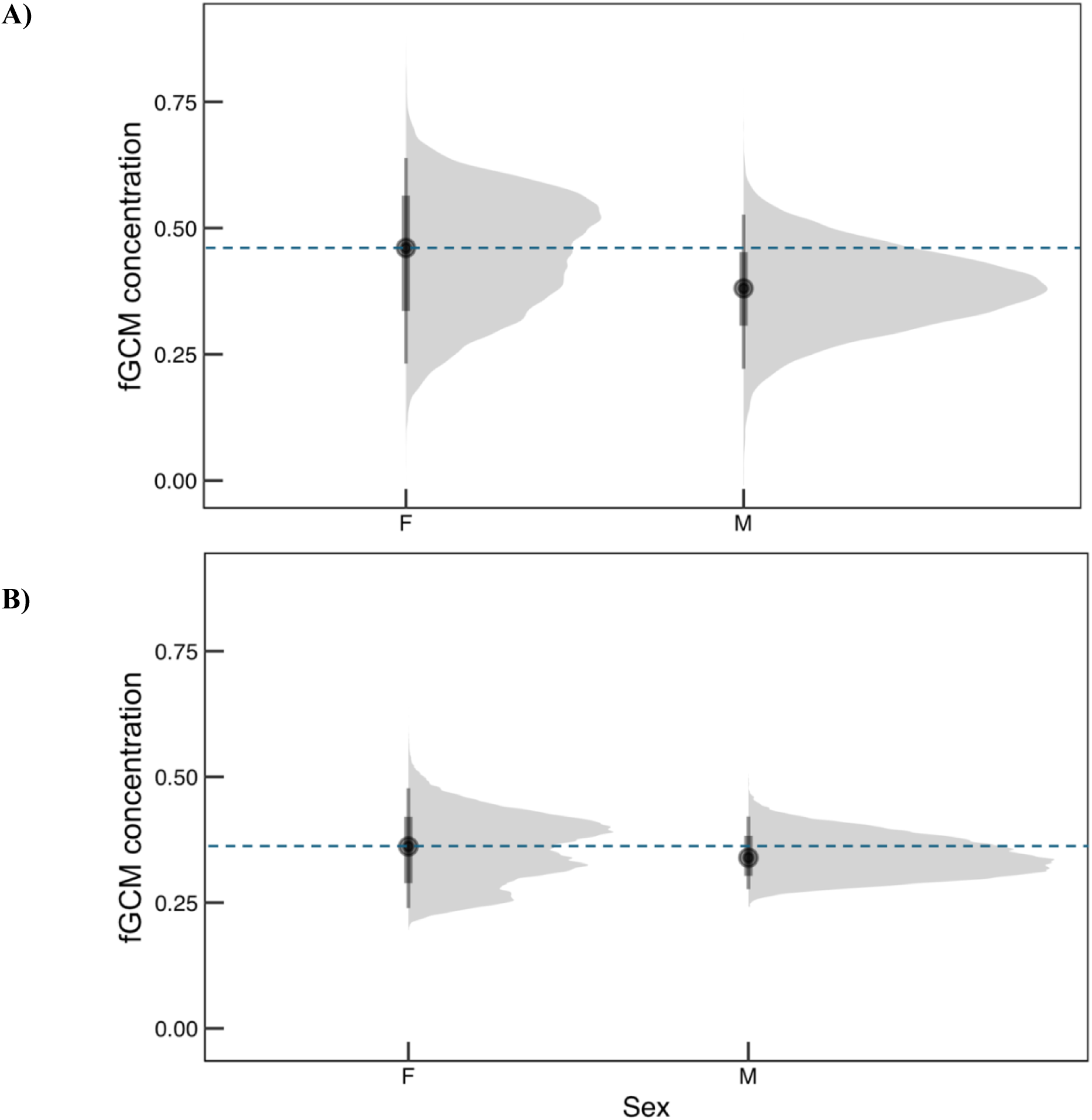
Posterior expectations displaying the relationship between sex and faecal glucocorticoid metabolite concentrations (fGCMs) within a A) disturbance period following helicopter use within the biophysical environment (Model 5), and a B) non-disturbance period within the biophysical environment (Model 6). fGCM concentrations are in μg/g dry weight. Density plots indicate the range of fGCM levels predicted by the models for females and males. The black dot represents the mean, and the dark bar represents the 95% credible intervals. We set our reference variable to female elephants and have a dashed line through the mean of these reference variables. F: female; M: male.

**Figure 4.**
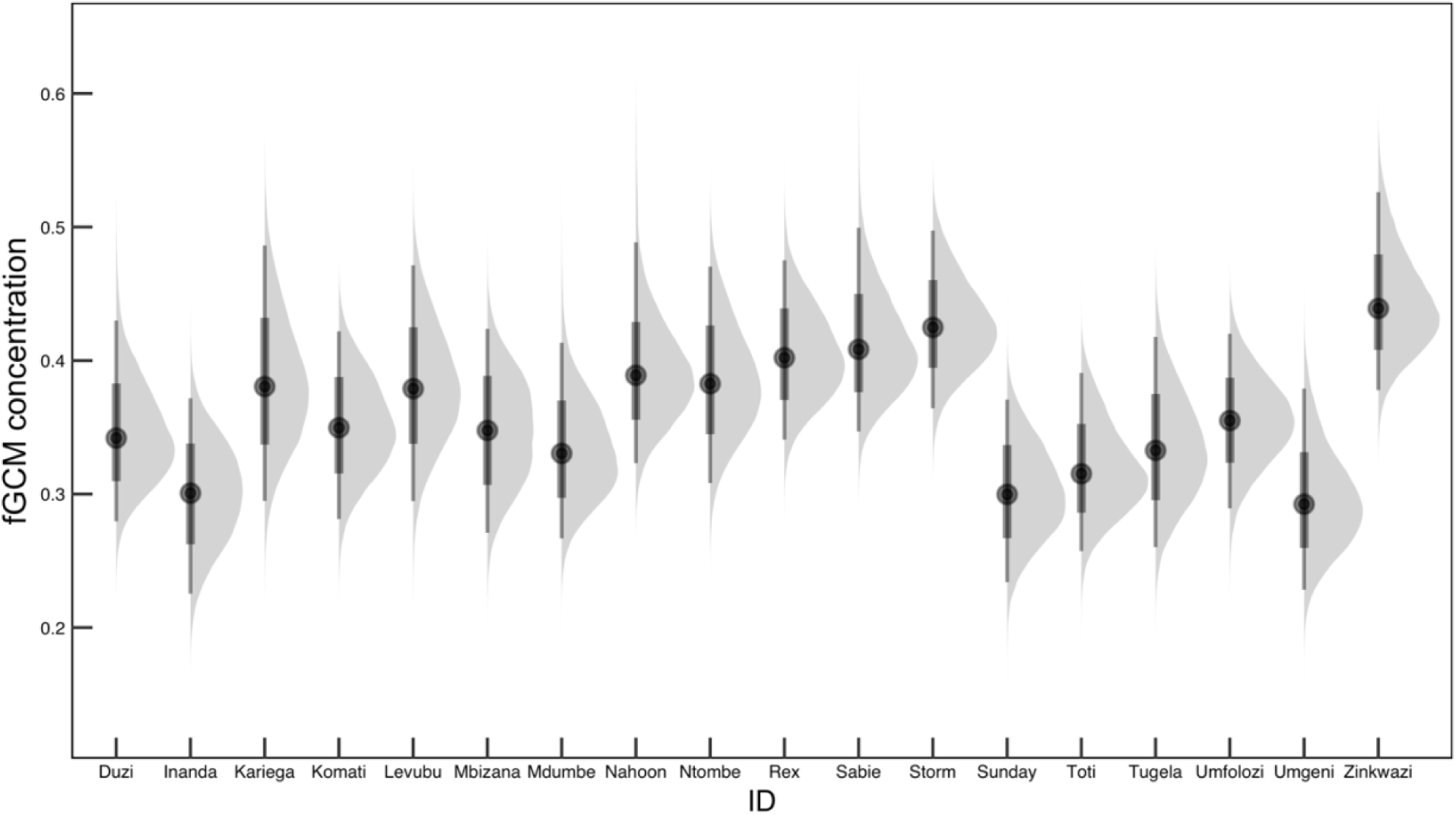
Posterior expectations which display the relationship between individual elephant and faecal glucocorticoid metabolite concentrations (fGCMs). fGCM concentrations are in μg/g dry weight. Density plots indicate the range of fGCM levels predicted by Model 1 for each individual. The black dot represents the mean predicted fGCM concentration. The thick vertical line represents the 66% credible interval, while the thin vertical line represents the 95% credible interval around the posterior estimate.

## 4 Discussion

The social environment and biophysical environment act together to shape individual responses to perceived stressors, with each either compounding the physiological response from the other or mitigating its effects and acting as a “buffer” (Dantzer et al., 2014; MacLeod et al., 2023). The social and biophysical environment must therefore be examined in concert to determine how they are influencing the stress response (Crespi et al., 2013; MacLeod et al., 2023). Consistent with the social buffering hypothesis (Cohen & Wills, 1985; Kikusui et al., 2006), dominance rank mediated fGCM concentrations differently across biophysical contexts, supporting our prediction that socially favourable individuals (i.e., higher ranking-individuals) would exhibit comparatively lower fGCM concentrations following disturbance periods, whereas lower-ranking individuals would exhibit comparatively lower fGCM concentrations during non-disturbance periods. Although dominance rank alone did not predict fGCM levels in this population, rank-related patterns shifted with biophysical context (Figure 2A), following the helicopter disturbance period, where lower-ranking individuals showed stronger associations with higher fGCMs compared to high-ranking individuals. The opposite was true during non-disturbance periods within the biophysical environment, where lower-ranking individuals showed weaker associations with higher fGCM concentrations compared to high-ranking individuals. This pattern suggests dominance rank mediates fGCM concentrations differently depending on external conditions, rather than exerting a consistent effect on physiological stress (Bonier, Martin, et al., 2009).

Degree centrality, or the number of connections an individual has within the network, predicted fGCM concentrations in non-disturbance periods, but did not predict fGCM concentrations in disturbance periods following helicopter use. We also found individuals with fewer social connections exhibited higher fGCM concentrations in both non-disturbance periods, and disturbance periods following helicopter use compared to individuals with more social connections (Figure 2B), suggesting that socially connected individuals may experience greater physiological buffering. Taken together, these findings support the social buffering hypothesis, where more affiliative connections may reduce physiological stress. However, the absence of a detectable effect under disturbance should be interpreted cautiously, as the model including disturbance periods following helicopter use was based on a smaller sample size than the model for non-disturbance conditions. Perhaps with greater sampling during disturbance periods, social effects may become more apparent. In other species, e.g., chimpanzees, social support is known to reduce GCs during both stressful events and everyday non-stressful interactions (Wittig et al., 2016). The physiological impacts of social relationships have also been documented in elephants, where juveniles with more individuals within their age cohort have lower fGCM concentrations (Parker et al., 2022). Given that social connectedness predicted fGCM concentrations in non-disturbance periods, but not under disturbance periods following helicopter use, social affiliations may play a stronger role in buffering baseline physiological variation, compared to acute anthropogenic stressors.

Betweenness centrality, which measured the extent to which an individual lies between other individuals within the network, did not predict fGCM levels, and patterns did not shift with biophysical context (Figure 3C). However, individuals with high betweenness centrality scores tended to have higher fGCM compared to individuals with low betweenness centrality scores, suggesting that occupying brokerage positions may carry greater physiological demands. In many social systems, individuals with high betweenness centrality scores may benefit as access to information, resources, or social influence, are more likely to be transmitted through them; they serve as “brokers” that link otherwise weakly connected or disconnected subgroups and may mediate the interactions between them (Farine & Whitehead, 2015; Wey et al., 2008). However individuals occupying brokerage positions may also experience higher social demands, conflict, or pathogen exposure, which has physiological costs (Creel, 2005; Evans et al., 2021; Muller et al., 2021). Individuals moving frequently among subgroups may repeatedly negotiate social relationships and maintain or establish such relationships with multiple known or unknown partners. Such repeated social interactions could increase physiological demands, particularly for dispersing males. Additionally, occupying a brokerage position and having multiple, seemingly redundant ties may act as a bet-hedging strategy by buffering individuals against the loss of key partners or changes in group composition (Firth et al., 2017; Pinter-Wollman et al., 2013). For instance, in bottlenose dolphins, a fission–fusion species, central individuals with high-betweenness centrality scores maintain social cohesion and bridge otherwise weakly connected subgroups (Lusseau, 2007; Lusseau & Newman, 2004). Consistent with this interpretation, Mdumbe, a young adult male in our study with the highest betweenness centrality score, regularly moved between individuals and subgroups (personal observations). Such behaviour likely reflects elevated social demands during the transition from natal family membership to participation in fluid all-male herds, when males must navigate changing relationships, fission-fusion dynamics, and emerging independence (Chiyo et al., 2011; Murphy et al., 2020; Vidya & Sukumar, 2005).

Contrary to H1 P2, neither individual aggressive interactions nor intergroup aggressive interactions predicted fGCM scores. One possibility for this finding is that the dominance hierarchy in this population is relatively stable, given the lack of emigration or immigration of new individuals, and the hierarchy would not need to be tested as frequently. An alternate explanation is that most females originated from the same founding family group and would not need to establish or test dominance relationships with new individuals compared to populations that fuse and separate from other family groups over time. Additionally, the aggression metrics we used, following Wittemyer & Getz, (2007), capture overt gestural interactions, but do not reflect more subtle or imperceptible forms of social communication such as vocal signalling or olfactory communication (O’Connell-Rodwell et al., 2011; Stoeger & Baotic, 2016; von Dürckheim et al., 2022; Eleuteri et al., 2025).

We also found no evidence that the natural biophysical indicators we examined, such as weather, water availability, or season, influenced fGCM concentrations. These findings align with previous work in this population showing no effect of seasonal or climatic variables on body condition, likely due to relatively stable principal forage species available within the reserve (Funk, Brown, et al., 2025). Such stability suggests that elephants in this system do not experience energetic limitations severe enough to elevate fGCM levels. In contrast, helicopter use strongly predicted increases in fGCM concentrations, indicating that anthropogenic disturbance represents the primary biophysical stressor in this population, and highlighting the measurable, physiological consequences of routinely used management interventions in this species.

Differences in fGCM concentrations were also evident between sexes. Females consistently exhibited higher fGCM concentrations across contexts than males, and the effect was stronger following helicopter use (Figure 3). Helicopters are used in the administration of porcine zona pellucida (pZP) immunocontraception to females of this population, and although the entire group is exposed to the disturbance, the females are specifically targeted and darted from a low-flying helicopter to administer fertility control. Although the pZP vaccine is non-hormonal and direct endocrine effects cannot be assumed, repeated and prolonged fertility control administration (which is needed for long-term efficacy) may impose physiological costs beyond the acute disturbance itself and merits further investigation (Funk, Ferraro, et al., 2025). Notably, these sex discrepancies were apparent in our population despite the absence of energetically costly reproductive states, i.e., no pregnancies or lactating cows. Additionally, in a previous study of this population within the same study period, adult females had significantly better body condition over time compared to all other age-sex categories (Funk, Brown, et al., 2025). Elevated fGCM concentrations in females therefore do not appear to reflect energetic limitation in this system. Instead, these patterns suggest that repeated disturbance associated with helicopter use may elevate glucocorticoid output even when individuals maintain good body condition. High among-individual variation in all models further reinforces the need for caution when interpreting GC measures without repeated sampling and detailed individual level context. Replication across additional fenced populations is also required to determine the generalizability of our findings, especially given the small sample size in our study.

## 5 Conclusion/Implications for Management and Conservation

Our results demonstrate how social and biophysical environments jointly shape physiological responses to stressors in a managed elephant population. In this population, alterations in individual stress-related fGCM concentrations reflect an interaction between anthropogenic disturbance, sex, and social network measures rather than natural biophysical stressors such as severe weather, low water availability, and seasonal change. Following helicopter disturbance, low-ranking individuals showed stronger associations with higher fGCM concentrations, suggesting greater physiological responses to anthropogenic disturbance compared to high-ranking individuals. The opposite was true during low disturbance periods. Such context dependence suggests dominance rank may buffer fGCM levels from rising when conditions are potentially stressful. Social connectedness further contributed to variation in stress-related physiological responses, where individuals with higher degree centrality had comparatively lower fGCM concentrations, suggesting stronger physiological buffering in more socially connected individuals. However, this effect was only present under non-disturbance periods, with no detectable effects during disturbance periods following helicopter use. This pattern suggests that social connections shape baseline physiological variation, but may be insufficient to buffer against acute anthropogenic stressors.

Our finding that social factors can mediate physiological responses to disturbance has direct implications for conservation and management. Notably, helicopter use, which is commonly used in elephant management interventions across Africa, can elicit measurable physiological stress responses, particularly in socially or demographically vulnerable individuals, even in well-monitored reserves. Because these effects are mediated by social factors, management interventions that disrupt social networks or disproportionately affect specific individuals must be approached with caution, given the potential for cascading consequences for population-level physiological effects and social buffering. Monitoring fGCM concentrations alongside social factors can help identify when management interventions impose disproportionate costs, and on which individuals. Integrating endocrine and social network approaches with long-term individual behavioural monitoring provides a powerful framework for evaluating animal welfare and resilience of managed wildlife populations.

## Conflict of interest

None declared

## Acknowledgments

Funding was provided by Natural Sciences and Research Council of Canada grants to K.R.F. and E.VW. K.R.F. is also supported by graduate funding through the Memorial University of Newfoundland. We are sincerely grateful to Kitty, Richard, and Danny Viljoen who generously provided field equipment, transportation, accommodation, and logistical support. We thank the Wildlife Evolutionary Ecology Lab for useful feedback and discussions. We greatly appreciate the comments provided by two anonymous referees which greatly improved the manuscript. Thank you to the owners and managers of Asante Sana Private Game Reserve for permission to work on the property.

## Author contributions CRediT

K.R.F: Conceptualization, Formal analysis, Funding acquisition, Investigation, Methodology, Writing – original draft, Writing – review and editing

L.N: Conceptualization, Writing – original draft, Writing – review and editing

L.R.B: Supervision, Writing – review and editing

A.G: Investigation, Data curation, Writing – review and editing

E.VW: Conceptualization, Funding acquisition, Writing – review and editing, Supervision

## Data Availability

Data and code supporting the findings of this study are available through the following link: https://github.com/krfunk/Social-buffering-karoo-elephants

## Appendix

**Table A1.** Operational definitions of overt dominance interactions.

*Operational definitions of overt dominance interactions*
| Interaction class | Interaction observed | Operational definition |
| --- | --- | --- |
| Physical | Chases | A runs in the direction of B, B then runs away from A |
|  | Pokes | A contacts B with a tusk, B then moves away from A |
|  | Pushes | A contacts B using a body part other than a tusk, B then moves away from A |
| Nonphysical | Supplants | A moves without running directly towards B, typically with ears flared, B then moves away from A, A takes place of B |
|  | Displaces | A moves without running directly towards B, typically with ears flared, B then moves away from A |

Operational definitions are adapted from Wittemyer & Getz (2007).

**Table A2.** Posterior Estimates of fGCM Scores from Model 1.

*Posterior Estimates of fGCM Scores from Model 1*
| Effect | Parameter | Estimate | S.E. | l-95% CI | u-95% CI | Rhat | Bulk ESS | Tail ESS | PD |
| --- | --- | --- | --- | --- | --- | --- | --- | --- | --- |
| Population-Level Effects | Intercept | -2.60 | 0.09 | -2.77 | -2.42 | 1.00 | 596 | 999 | 100% |
|  | Rainfall | 0.01 | 0.01 | 0.00 | 0.02 | 1.00 | 5090 | 3318 | **97.94%** |
|  | Windspeed average | -0.00 | 0.01 | -0.02 | 0.01 | 1.00 | 3959 | 3176 | 64.98% |
|  | Temperature average | 0.02 | 0.01 | -0.01 | 0.04 | 1.00 | 3027 | 2492 | 87.90% |
|  | Water available in dams | 0.02 | 0.71 | -1.34 | 1.41 | 1.00 | 2926 | 2879 | 50.22% |
|  | Specific season WS1 | -0.03 | 0.02 | -0.07 | 0.01 | 1.00 | 3592 | 3592 | 91.62% |
|  | Specific season WS2 | 0.01 | 0.71 | -1.37 | 1.36 | 1.00 | 2923 | 2860 | 51.52% |
|  | Low/high temperature difference | -0.01 | 0.01 | -0.03 | -0.00 | 1.00 | 4011 | 3459 | **98.06%** |
|  | Atmospheric pressure | 0.00 | 0.01 | -0.01 | 0.02 | 1.00 | 4528 | 3310 | 75.04% |
| Group-Level Effects | sd(individual ID) | 0.35 | 0.07 | 0.25 | 0.51 | 1.00 | 849 | 1313 | -- |

Seasons are relative to the dry season. “Low/high temperature difference” refers to the difference between the minimum and maximum temperature recorded each day. fGCM: faecal glucocorticoid metabolite concentrations. SE: Standard error of the estimate, CI: credible interval, Bulk ESS: bulk effective sample size estimate, Tail ESS: tail effective sample size estimate, PD: probability of direction. $N = 372$ . $\text{Loo } R^2 = 0.177$ .

**Table A3.**
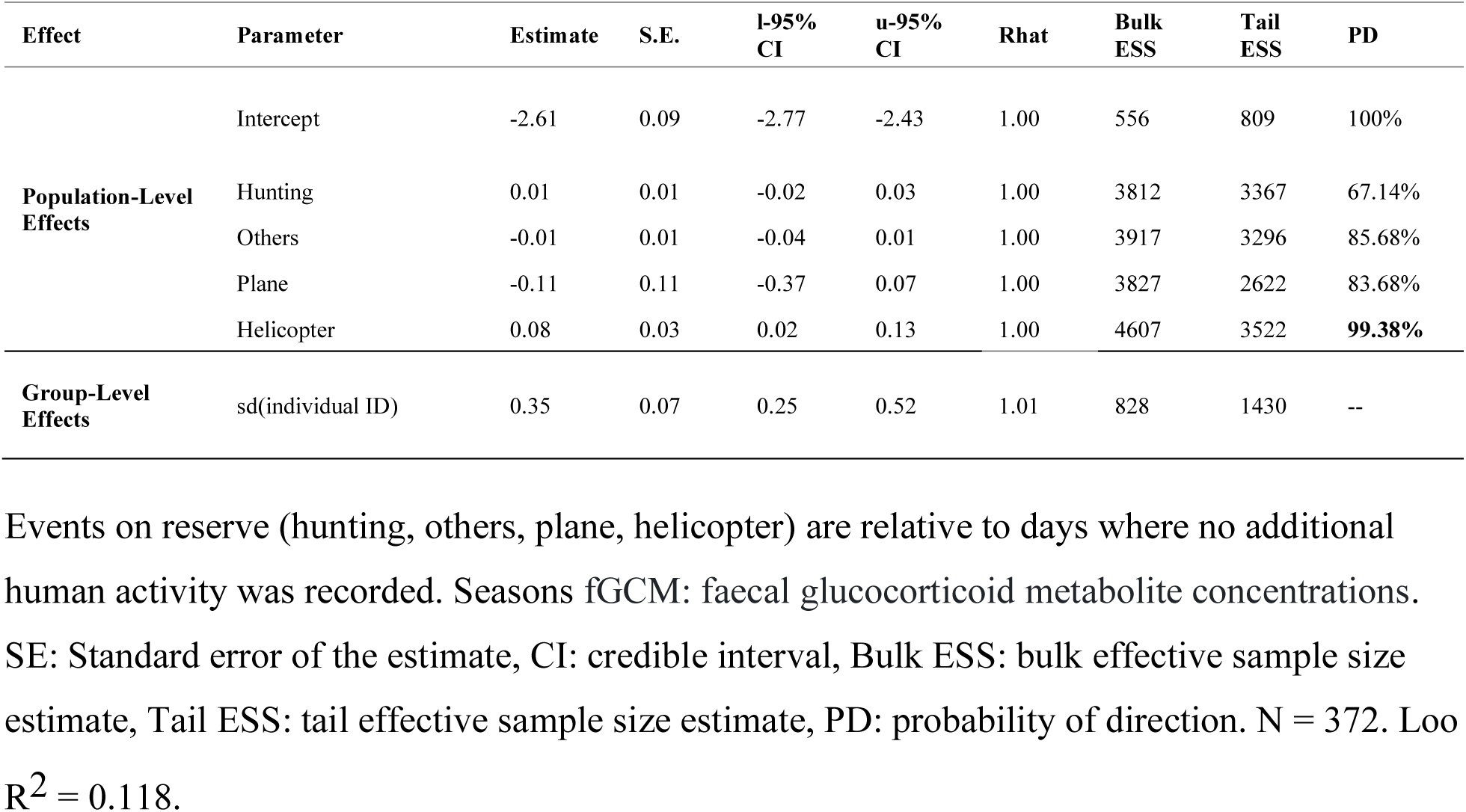
Posterior Estimates of fGCM Scores from Model 2.

**Table A4.**
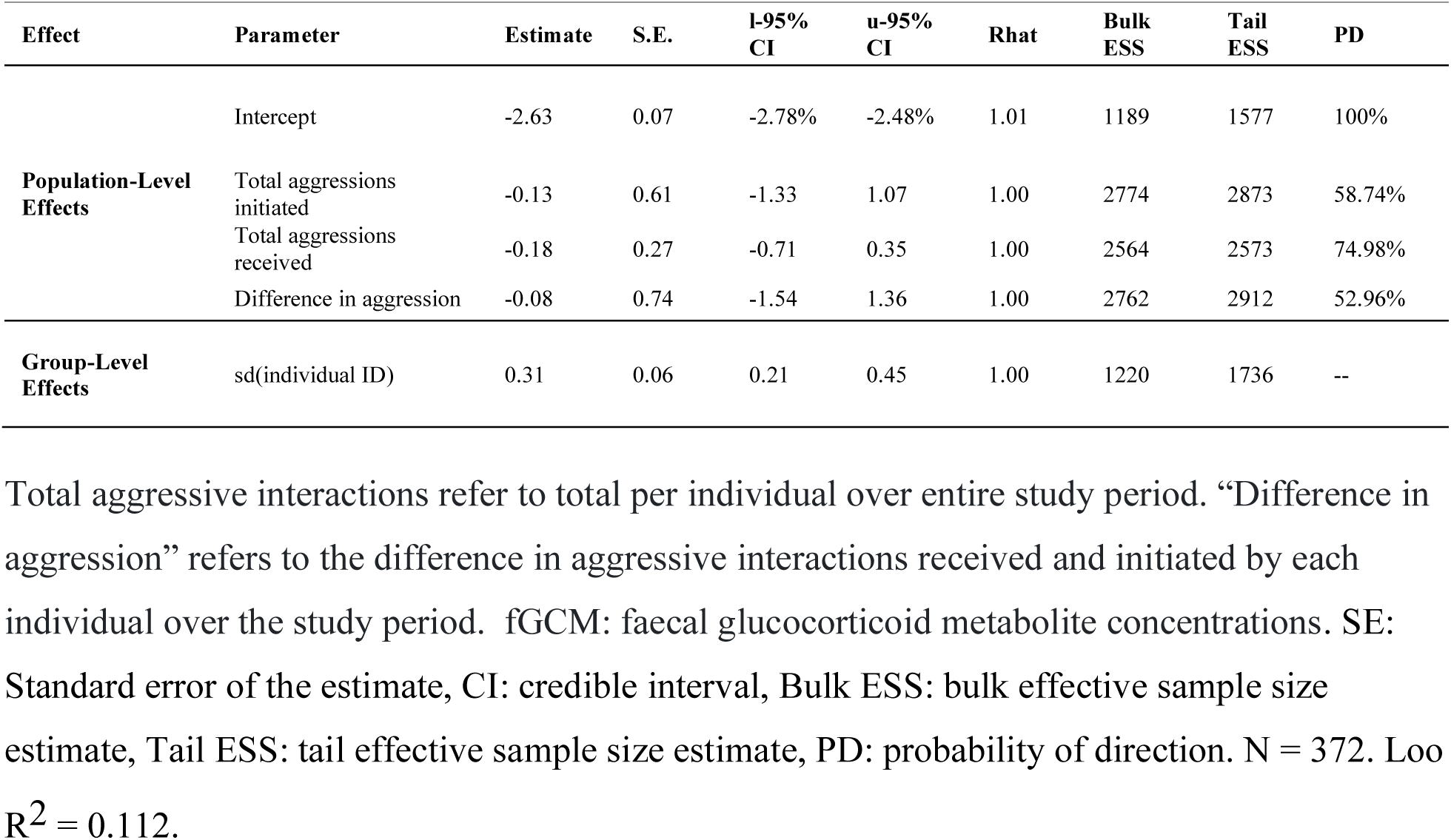
Posterior Estimates of fGCM Scores from Model 3.

**Table A5.**
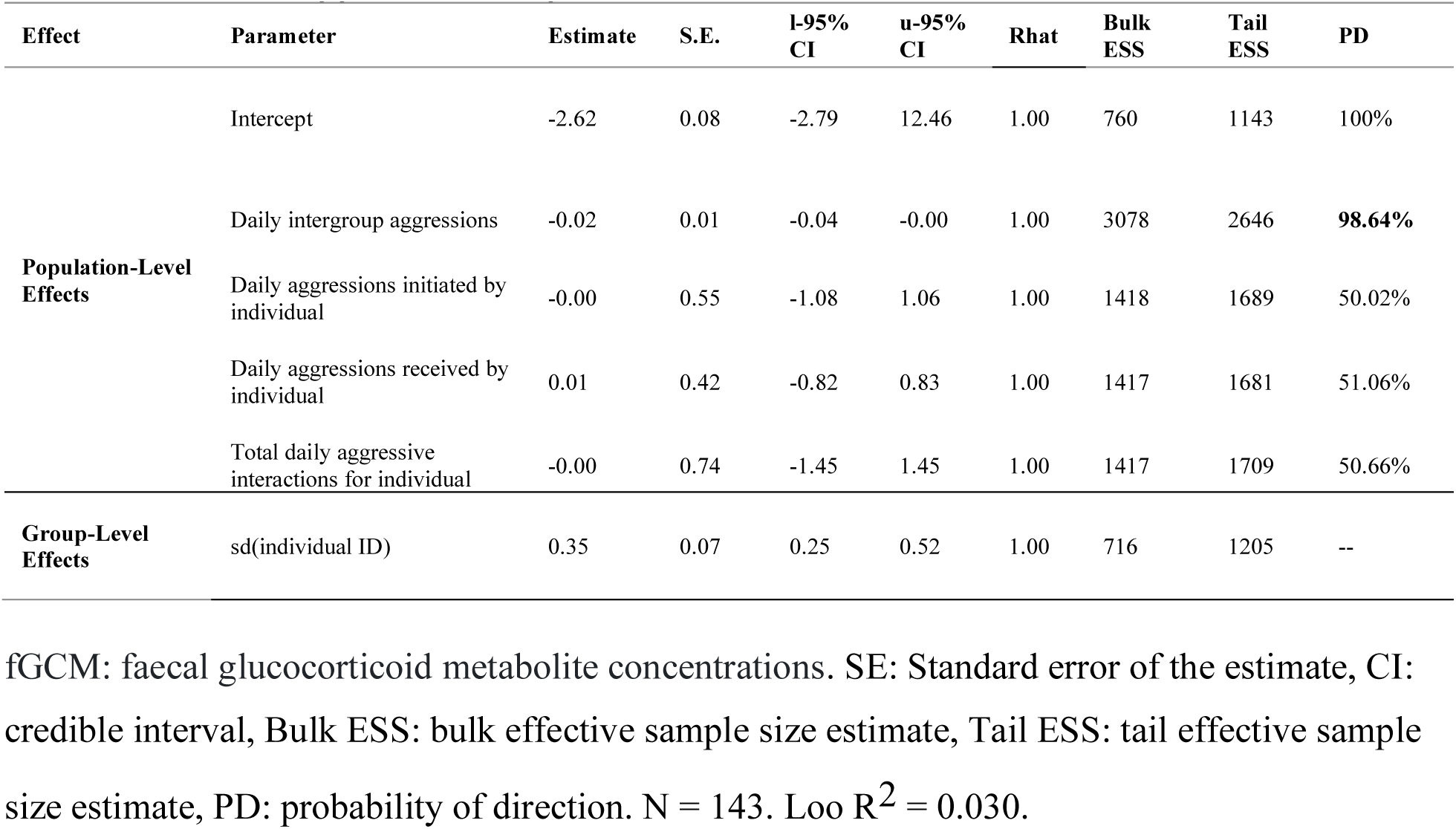
Posterior Estimates of fGCM Scores from Model 4.

**Table A6.**
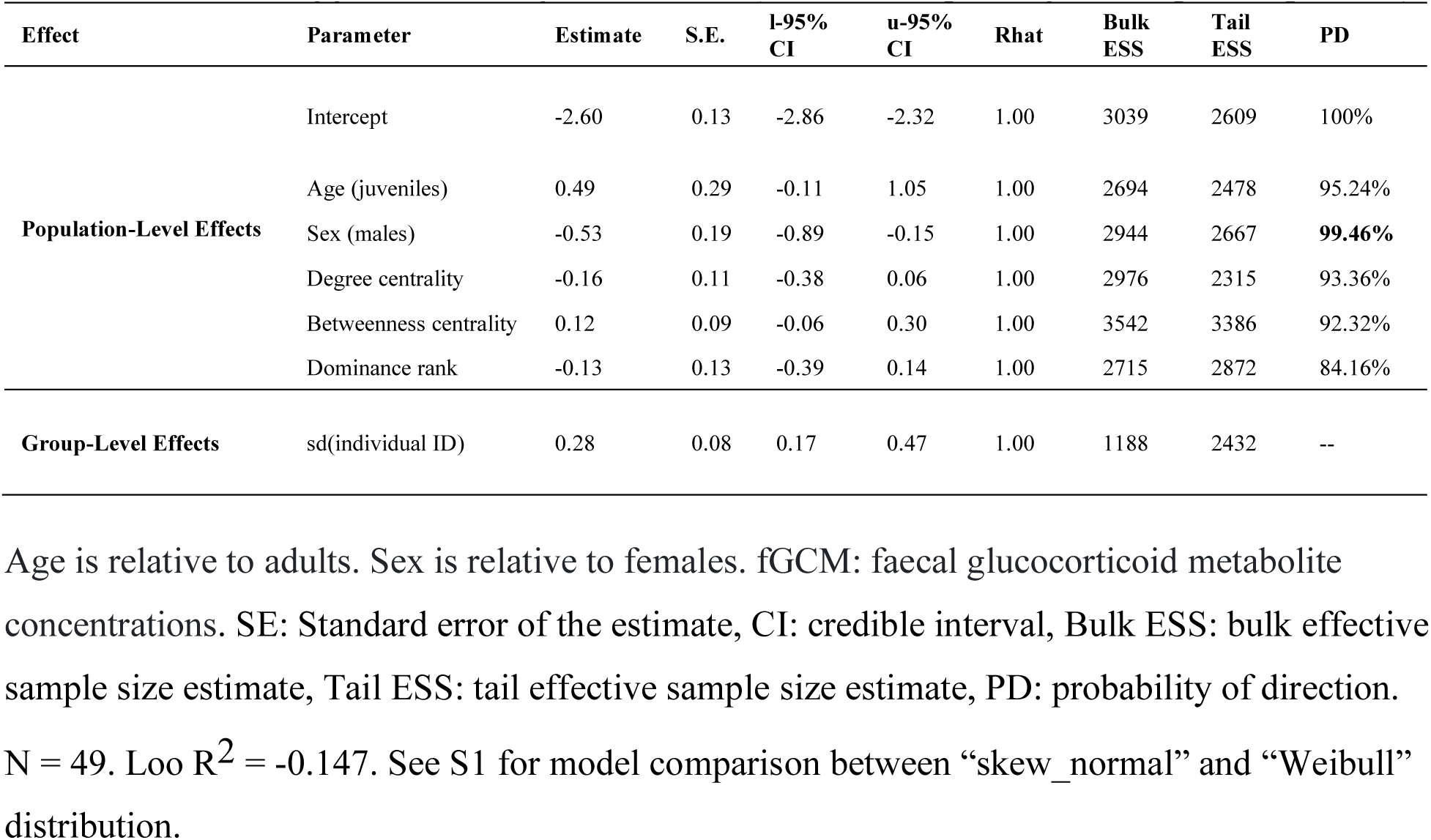
Posterior Estimates of fGCM Scores from Model 5 (disturbance period following helicopter use)

**Table A7.** Posterior Estimates of fGCM Scores from Model 6 (non-disturbance period)

| Effect | Parameter | Estimate | S.E. | l-95%<br>CI | u-95%<br>CI | Rhat | Bulk<br>ESS | Tail<br>ESS | PD |
| --- | --- | --- | --- | --- | --- | --- | --- | --- | --- |
| **Population-Level Effects** | Intercept | -2.61 | 0.10 | -2.80 | -2.42 | 1.00 | 1948 | 2549 | 100% |
|  | Age (juveniles) | 0.37 | 0.20 | -0.04 | 0.76 | 1.00 | 2766 | 2461 | 96.34<br>% |
|  | Sex (males) | -0.50 | 0.17 | -0.82 | -0.16 | 1.00 | 3228 | 3157 | **99.64**<br>% |
|  | Degree centrality | -0.18 | 0.08 | -0.33 | -0.01 | 1.00 | 3338 | 2744 | **98.26**<br>% |
|  | Betweenness centrality | 0.13 | 0.07 | -0.01 | 0.28 | 1.00 | 3625 | 2716 | 96.42<br>% |
|  | Dominance rank | 0.11 | 0.10 | -0.10 | 0.32 | 1.00 | 2872 | 2649 | 86.70<br>% |
| **Group-Level Effects** | sd(individual ID) | 0.24 | 0.06 | 0.16 | 0.39 | 1.00 | 2872 | 2649 | -- |

Age is relative to adults. Sex is relative to females. fGCM: faecal glucocorticoid metabolite concentrations. SE: Standard error of the estimate, CI: credible interval, Bulk ESS: bulk effective sample size estimate, Tail ESS: tail effective sample size estimate, PD: probability of direction.
N = 322. Loo $R^2$ = 0.035.

## Notes

https://github.com/krfunk/Social-buffering-karoo-elephants

### Competing Interest Statement

The authors have declared no competing interest.

