## Supplementary materials for "Sociality shapes context-dependent stress responses to anthropogenic disturbance in African elephants"

**Supplementary Materials**
**Figure S1**

A)

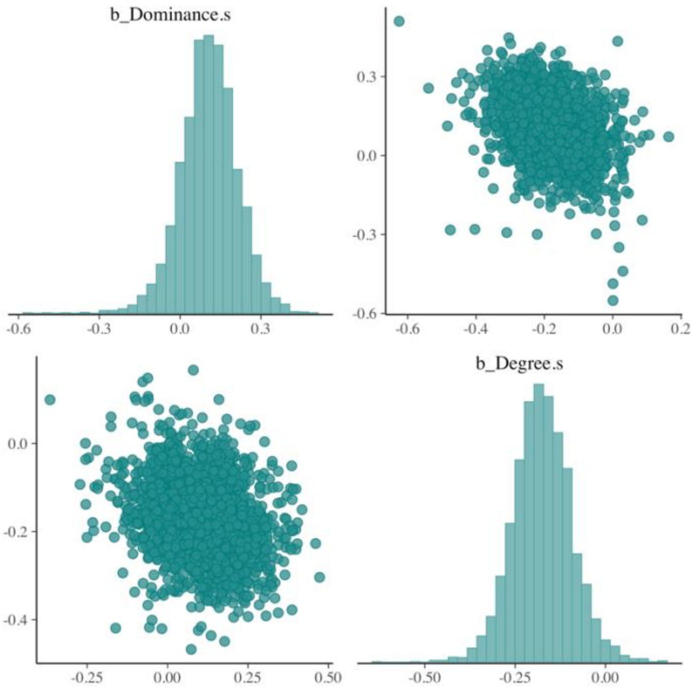


B)

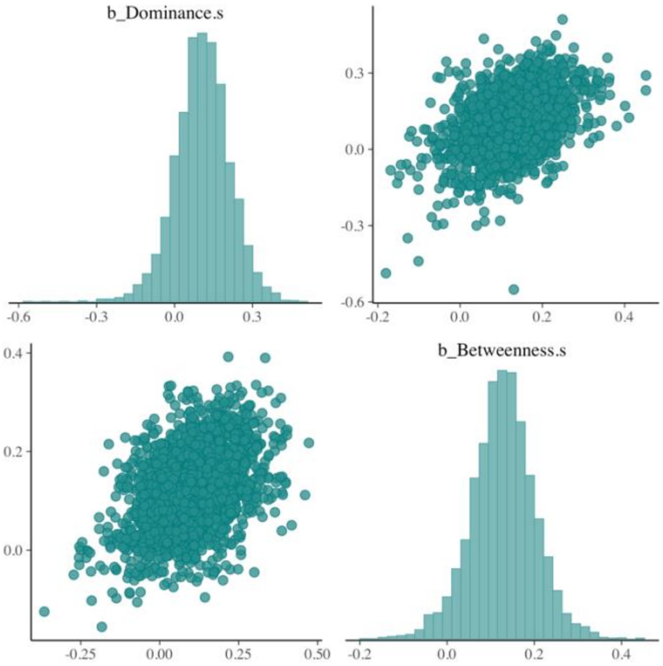

C)

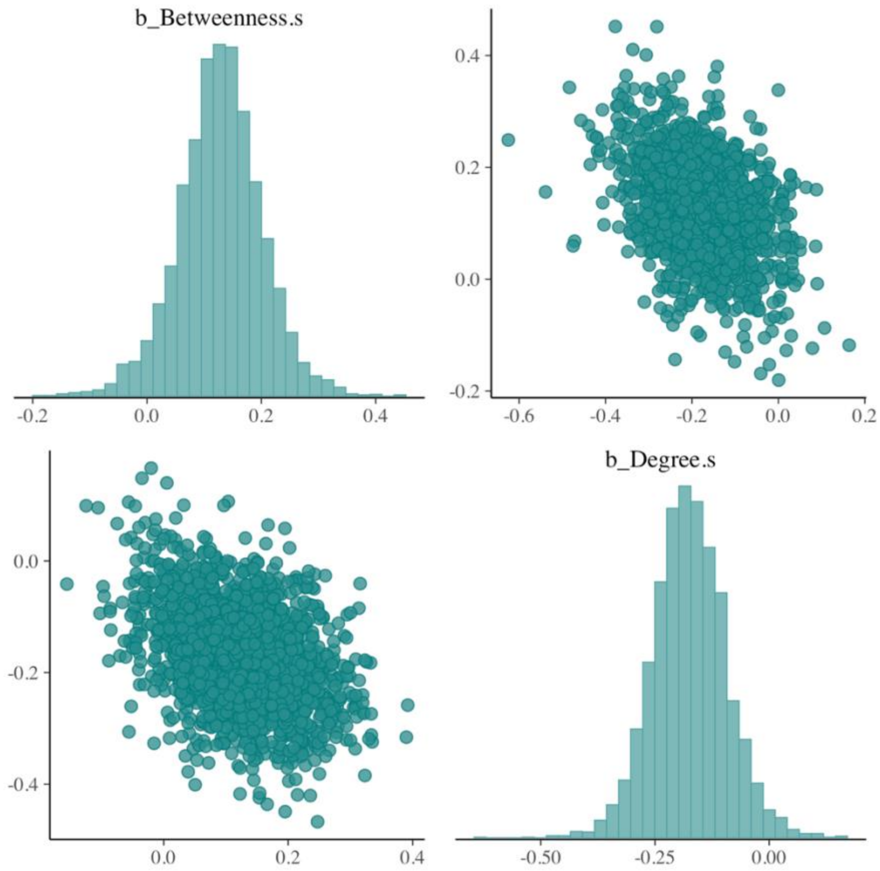


Pairwise plots comparing A) dominance and degree, *r*(320) = 0.487, B) dominance and betweenness, *r*(320) = -0.059, and C) betweenness and degree, *r*(320) = 0.161.

**Figure S2**

A)

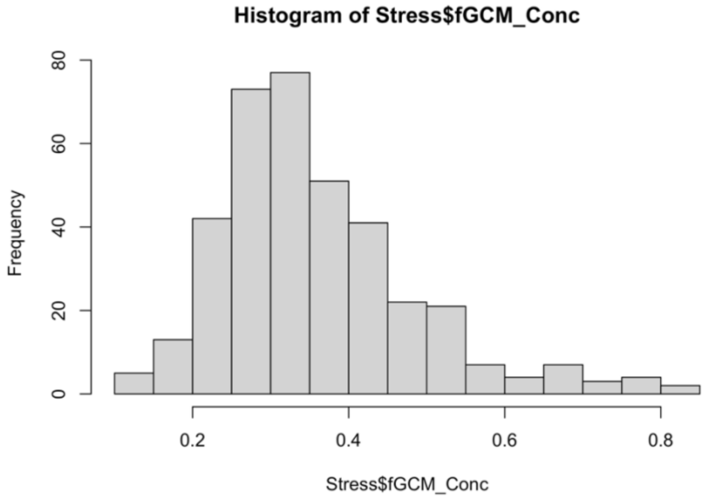


B)

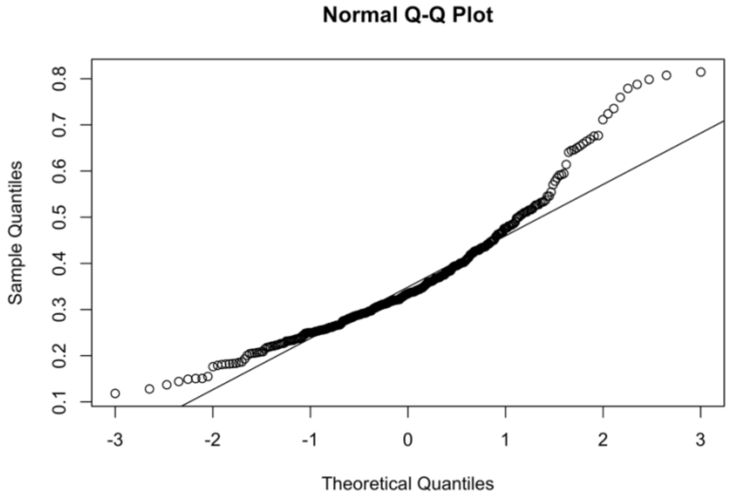


Histograms (A) of the faecal glucocorticoid metabolite concentrations and a (B) Q-Q plot of these data indicating a non-normal distribution.

**Figure S3**

| | pp_check | DHARMa nonparametric test |
|---|---|---|

Model 1

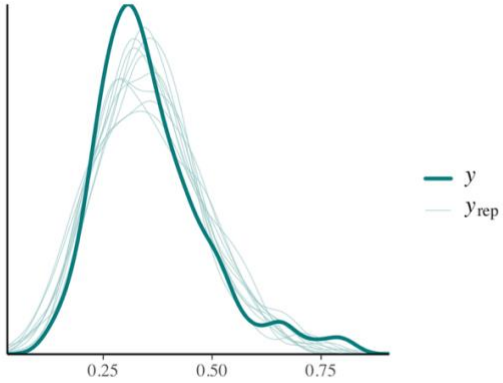


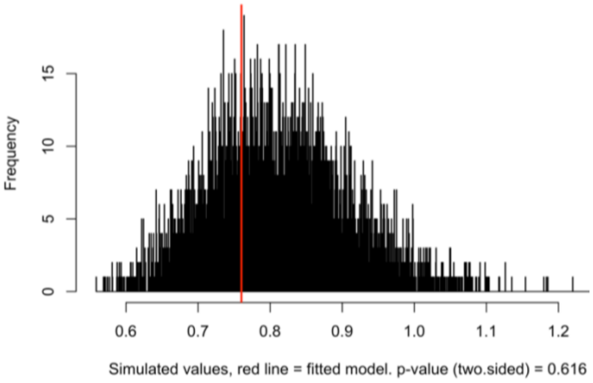


Model 2

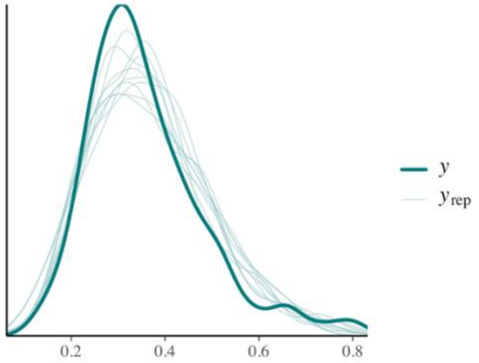


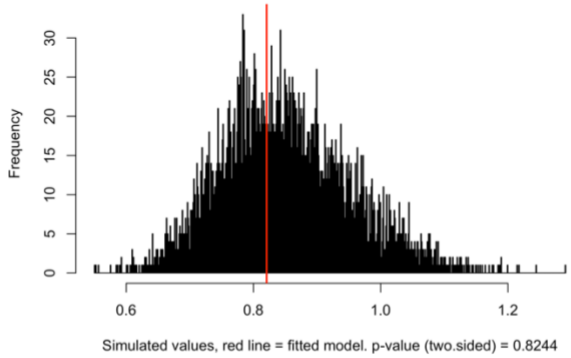


Model 3

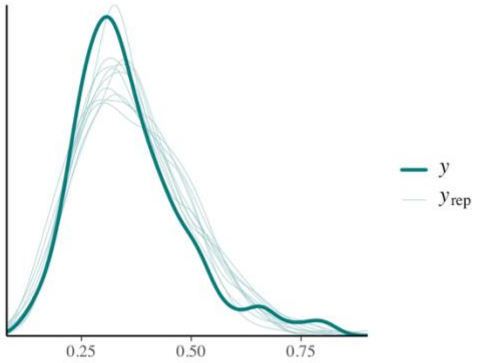


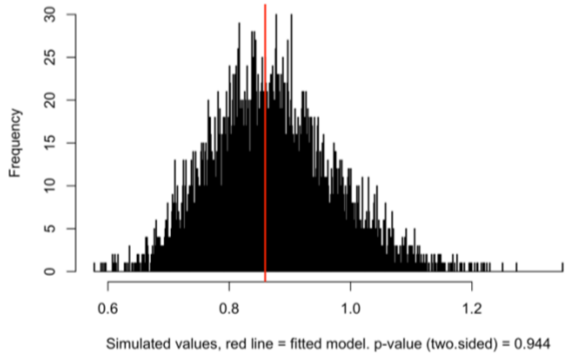


Model 4

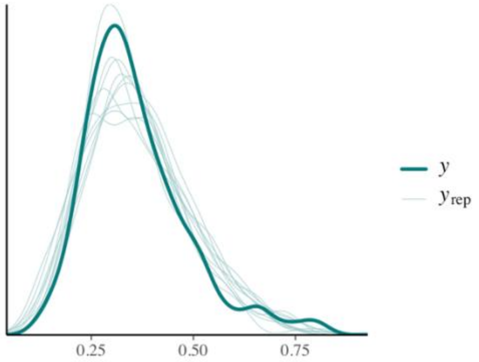


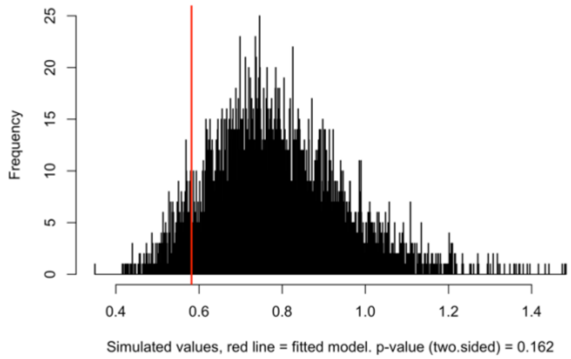

| | | |
|---|---|---|
| Model 5 (skew_normal distribution) | 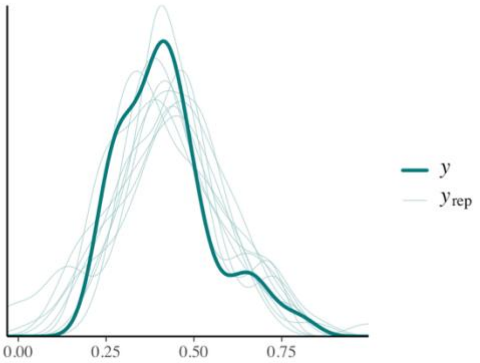 | 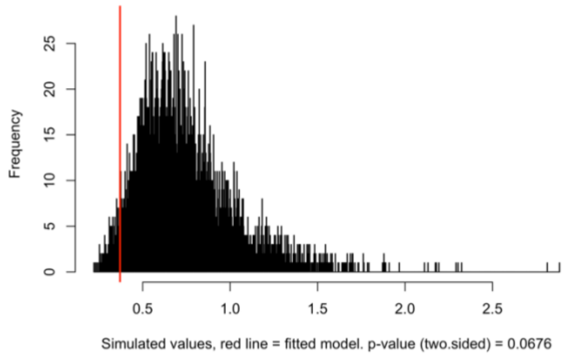 |
| Mode 5.2 (Weibull distribution) | 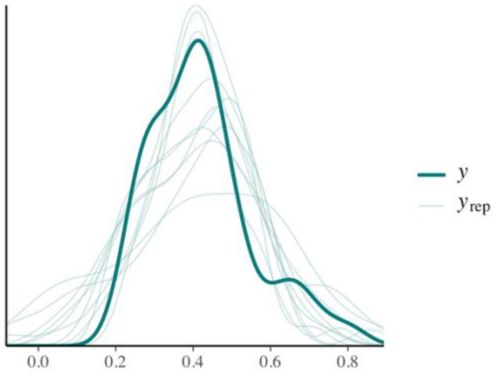 | 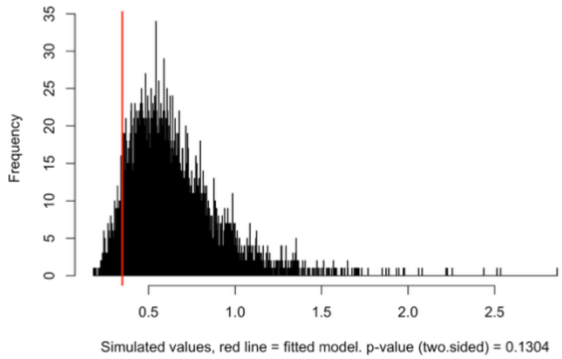 |
| Model 6 | 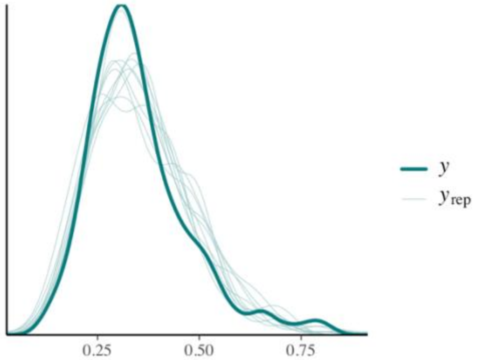 | 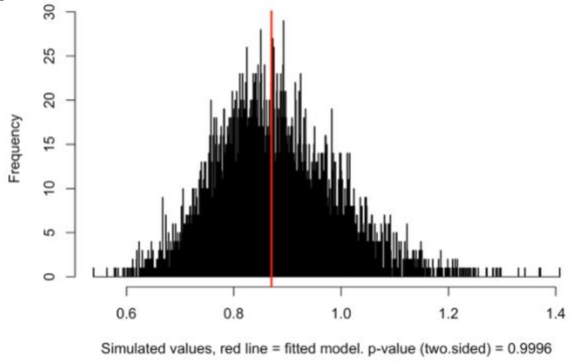 |



Using the ‘pp_check’ function to compare the observed outcome variable (faecal glucocorticoid metabolite concentrations) to datasets simulated from the posterior predictive distribution of the model. Where Y represents the actual representations from the model, where the YREP represents the possible model representations. The DHARMa nonparametric dispersion test via sd of residuals fitted vs. simulated. Both tests indicate no significant dispersion issues in any model.

**Table S1**

*Posterior Estimates of fGCM Scores from Model 5.2 with a Weibull Distribution*

| Effect | Parameter | Estimate | S.E. | l-95% CI | u-95% CI | Rhat | Bulk ESS | Tail ESS | PD |
|---|---|---|---|---|---|---|---|---|---|
| Population-Level Effects | Intercept | -3.88 | 0.20 | -4.28 | -3.47 | 1.00 | 3420 | 3135 | 100% |
| | Age (juveniles) | 0.61 | 0.42 | -.026 | 1.42 | 1.00 | 2608 | 2958 | 92.78% |
| | Sex (males) | -0.68 | 0.28 | -1.22 | -0.12 | 1.00 | 2477 | 2547 | **98.94%** |
| | Degree | -0.16 | 0.17 | -0.49 | 0.17 | 1.00 | 2638 | 2572 | 83.92% |
| | Betweenness | 0.14 | 0.14 | -0.15 | 0.41 | 1.00 | 2892 | 2766 | 84.82% |
| | Dominance rank | -0.21 | 0.19 | -0.59 | 0.19 | 1.00 | 2514 | 2542 | 87.30% |
| Group-Level Effects | sd(individual ID) | 0.43 | 0.12 | 0.25 | 0.72 | 1.00 | 1441 | 2276 | -- |

Age is relative to adults. Sex is relative to females. fGCM: faecal glucocorticoid metabolite concentrations. SE: Standard error of the estimate, CI: credible interval, Bulk ESS: bulk effective sample size estimate, Tail ESS: tail effective sample size estimate, PD: probability of direction. N = 49. Loo $R^2$ = 0.032.
